# Signed Reward Prediction Errors in the Ventral Striatum Drive Episodic Memory

**DOI:** 10.1101/2020.01.03.893578

**Authors:** Cristian Buc Calderon, Esther De Loof, Kate Ergo, Anna Snoeck, Carsten Nico Boehler, Tom Verguts

## Abstract

A growing body of behavioral evidence implicates reward prediction errors (RPEs) as a key factor in the acquisition of episodic memory. Yet, important neural predictions related to the role of RPE in declarative memory acquisition remain to be tested. Using a novel variable-choice task, we experimentally manipulated RPEs and found support for key predictions on the neural level with fMRI. Specifically, we demonstrate that trial-specific RPE responses in the ventral striatum (during learning) predict the strength of subsequent episodic memory (during recollection). Furthermore, functional connectivity between task-relevant processing areas (e.g., face-selective areas) and hippocampus, ventral tegmental area, and ventral striatum increased as a function of RPE value (during learning), suggesting a central role of these areas in episodic memory formation. Our results consolidate reinforcement learning theory and striatal RPEs as key operations subtending the formation of episodic memory.

## Introduction

When meeting a new person, being able to remember his/her name from a single encounter is essential. Referred to as episodic memory (Tulving, 1993), this information can for instance be recalled to strike up a conversation when running into that person later on.

Several studies have investigated the behavioral and neural mechanisms by which such one-shot episodic memories are learned. In particular, previous work identified an important role of reward. Compared to unrewarded contexts, items memorized within rewarding contexts are associated with better recognition performance in old-new item decisions (Shneyer & Mendelsohn, 2018). Neurally, this beneficial effect of reward on episodic memory has been ascribed to increased activity of the striatum. For instance, rewarded to-be-remembered items elicit stronger striatal activation when subsequently remembered (Adcock, Thangavel, Whitfield-Gabrieli, Knutson, & Gabrieli, 2006; for review see, Miendlarzewska, Bavelier, & Schwartz, 2016; Wittmann et al., 2005). Additionally, reward induces a gradual retroactive effect whereby items temporally closer to the reward are best remembered (Braun, Wimmer, & Shohamy, 2018).

However, learning based solely on reward has limited computational power. Reinforcement learning (RL) theory instead has highlighted the importance of reward prediction errors (RPEs) for learning (Sutton & Barto, 2018). RPEs arise when choice outcomes deviate from their predictions. They are implemented via dopaminergic activity bursts, stemming from midbrain nuclei (i.e. ventral tegmental area (VTA) and substantia nigra (SN)), and broadcast to ventral striatum (VS) and other cortical and subcortical areas (Watabe-Uchida, Eshel, & Uchida, 2017).

In line with RL theory, recent work suggests a crucial role of RPEs in episodic memory. Behaviorally, memory encoding improves linearly with better-than-expected rewards, called signed RPEs (SRPEs) (De Loof et al., 2018; Jang, Nassar, Dillon, & Frank, 2019). Neurally, high-beta and high-alpha oscillatory SRPE signatures mirror behavioral SRPE effects (Ergo, De Loof, Janssens, & Verguts, 2019). The behavioral SRPE effect on episodic memory also correlates with hippocampal activation and functional connectivity between hippocampus and striatum (Davidow, Foerde, Galván, & Shohamy, 2016). In addition, striatal activation is positively correlated with SRPEs and predicts choice confidence levels in a delayed recognition test (Pine, Sadeh, Ben-Yakov, Dudai, & Mendelsohn, 2018). Finally, RPEs elicited during the recognition phase of a previously learned item list, control the decision criterion for old-new item decisions (Scimeca, Katzman, & Badre, 2016).

Although significant evidence points towards an RL-based acquisition of episodic memory, key neural RL predictions have yet to be tested; we proceeded with the following steps. First, we attempted to replicate the behavioral SRPE effect on recognition memory, and the linear SRPE pattern in VS. Second, and crucially, we tested a trial-to-trial SRPE slope effect on subsequent episodic memory accuracy. Third, we investigated if functional connectivity between stimulus-processing areas and hippocampus, VTA, and VS was modulated by SRPE value. Finally, we correlated VS activation (across subjects) with recognition memory. We addressed these RL predictions on episodic memory encoding using fMRI.

## Results

### Recognition memory improves with learning-evoked SRPE

Thirty participants performed the variable-choice task in the MR scanner (Fig. 1A; see also Ergo et al., 2019). Each trial started with a fixation cross (0.5 sec), followed by a celebrity face in the top part of the screen, together with four village pseudo-names. After exploring the display for four seconds, either 1, 2 or 4 names were framed. The participant’s task was to guess which village name was associated with the celebrity face. Participants had to choose between the framed names, which was followed by choice feedback. By systematically varying the number of framed village names, we were able to manipulate the signed reward prediction error (SRPE) received at choice feedback, on each trial. We computed the SRPE as *r – p*, where *r* is the observed reward (1 and 0 for correct and incorrect guesses, respectively), and *p* is the probability of making a correct choice. This probability is 1, 0.5 or 0.25, respectively for the 1-, 2- or 4-frame conditions. Hence, SRPEs could take on the values −0.5, −0.25, 0, 0.5, 0.75. Upon completion of the variable-choice task, participants performed the memory test (outside the scanner). During this test, participants again observed the 70 faces alongside the same four competing village names (shuffled relative to their previous positions in the display during the variable-choice task). They had to select the village name previously associated with the presented face and were instructed to provide a certainty rating on their choice afterwards. We imposed no time restrictions on either task.

**Figure 1.**
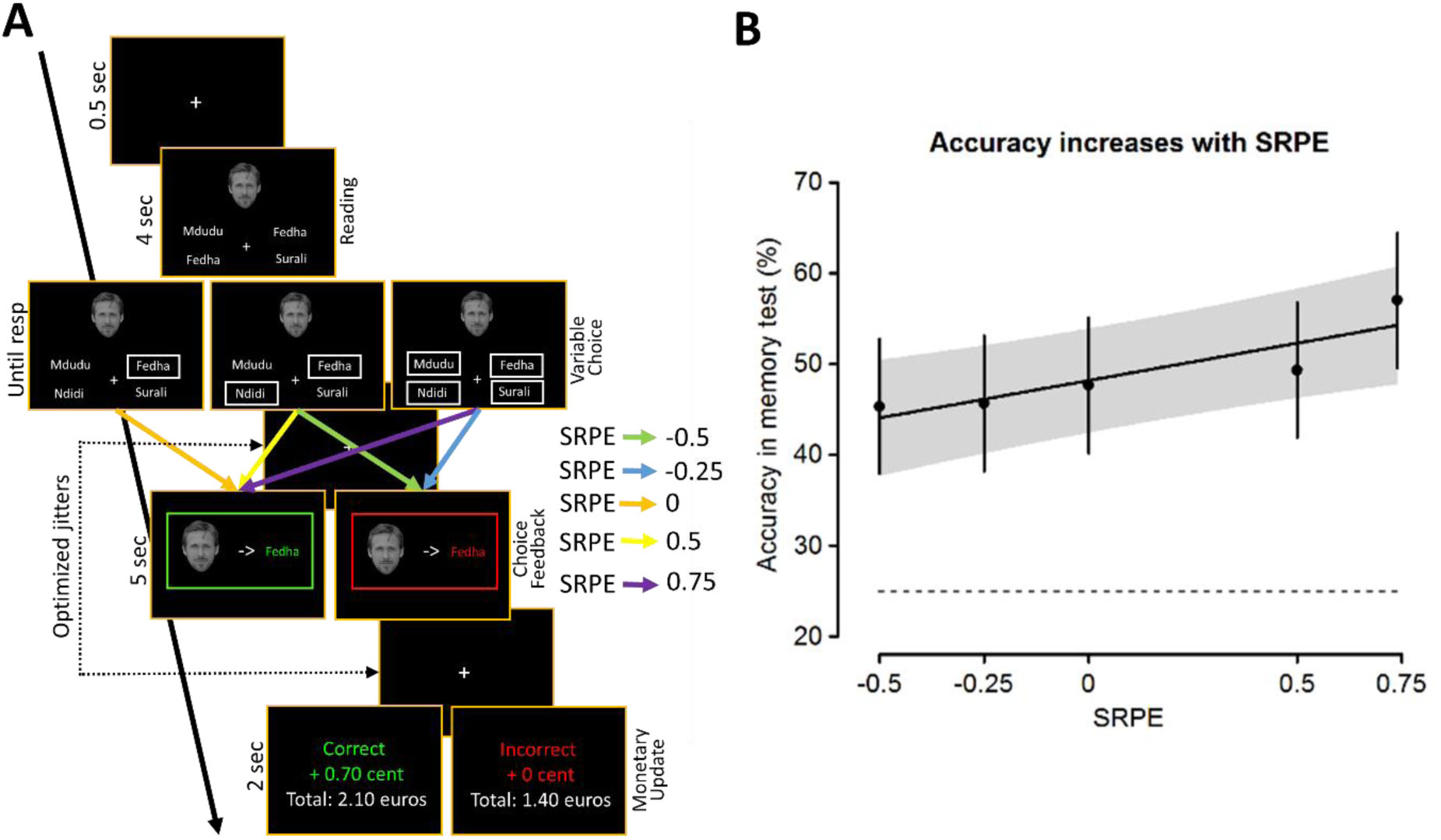
Behavioral paradigm and results. **A**. Experimental design. Following a fixation cross (0.5 sec), participants saw a face alongside four words (4 sec). Subsequently, either 1, 2 or 4 names were framed, indicating the options participants could choose from. Following a jittered interval with just a fixation cross, participants were shown the to-be-learned face-word association framed in green/red for correct/incorrect choices; this choice feedback evoked an SRPE of different levels (color-coded by the arrows between the variable-choice and choice-feedback events). Subsequently, they were shown a monetary update. **B**. Memory test behavioral results. A positive linear relationship between SRPE and accuracy on the subsequent memory test was observed; i.e., recognition accuracy increased as SRPE values increased. Error bars depict the 95% confidence interval; the dashed line indicates chance performance (25%).

As expected from previous work (De Loof et al., 2018; Jang et al., 2019) memory performance (i.e., recognition accuracy) increased linearly with SRPE, *χ*^*2*^(1, *N* = 30) = 11.18, *p* = 0.00083 (Fig. 1B). We observed a positive main effect of reward, *χ*^*2*^(1, *N* = 30) = 7.61, *p* = 0.0058. Importantly, we also observed a positive main effect of choice options number, *χ*^*2*^(1, *N* = 30) = 3.92, *p* = 0.048; thereby, suggesting that this linear increase was not due to a mere reward effect.

### Ventral striatum encodes SRPE

We next turn to fMRI data. All fMRI results (summarized in Table 1 and 2) are family-wise error (FWE) cluster-corrected; see Table 1 for contrast-specific voxel-wise thresholds.

**Table 1.**
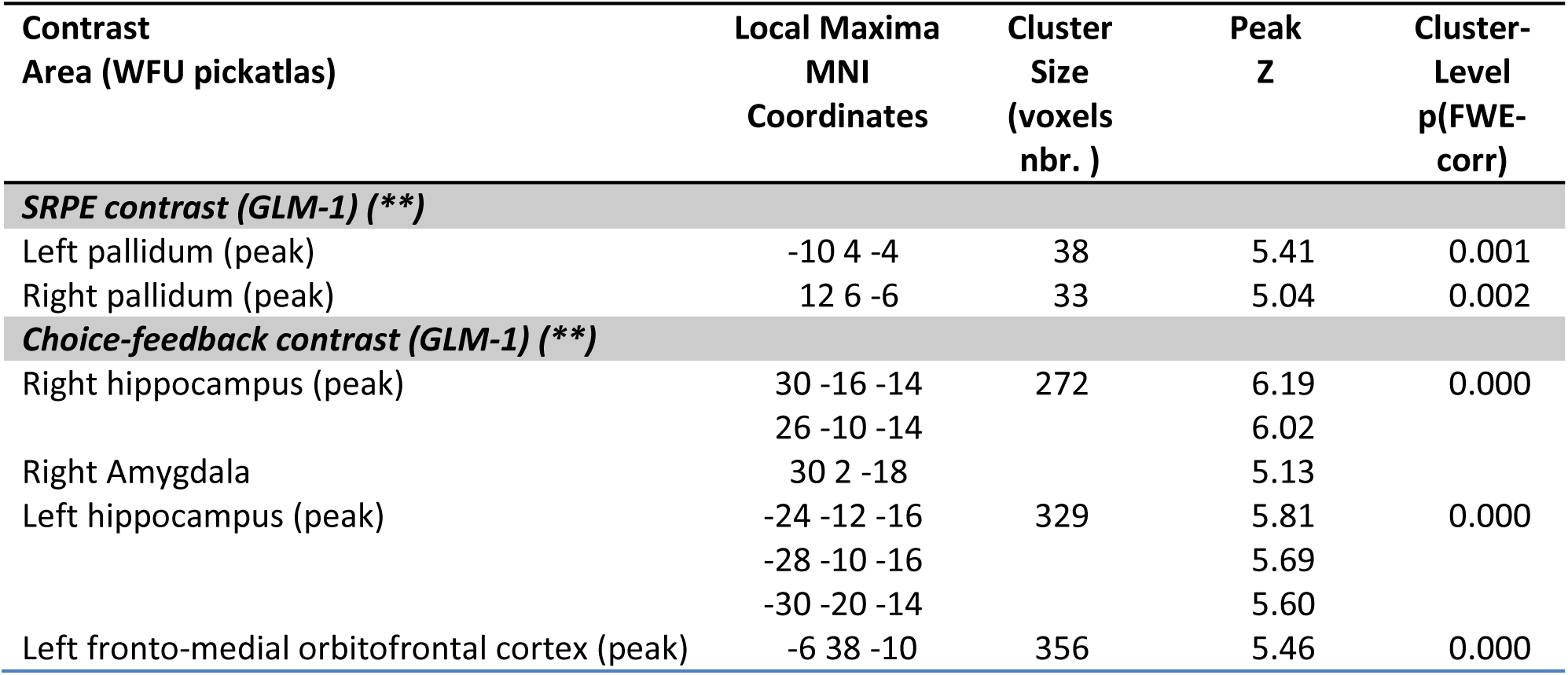
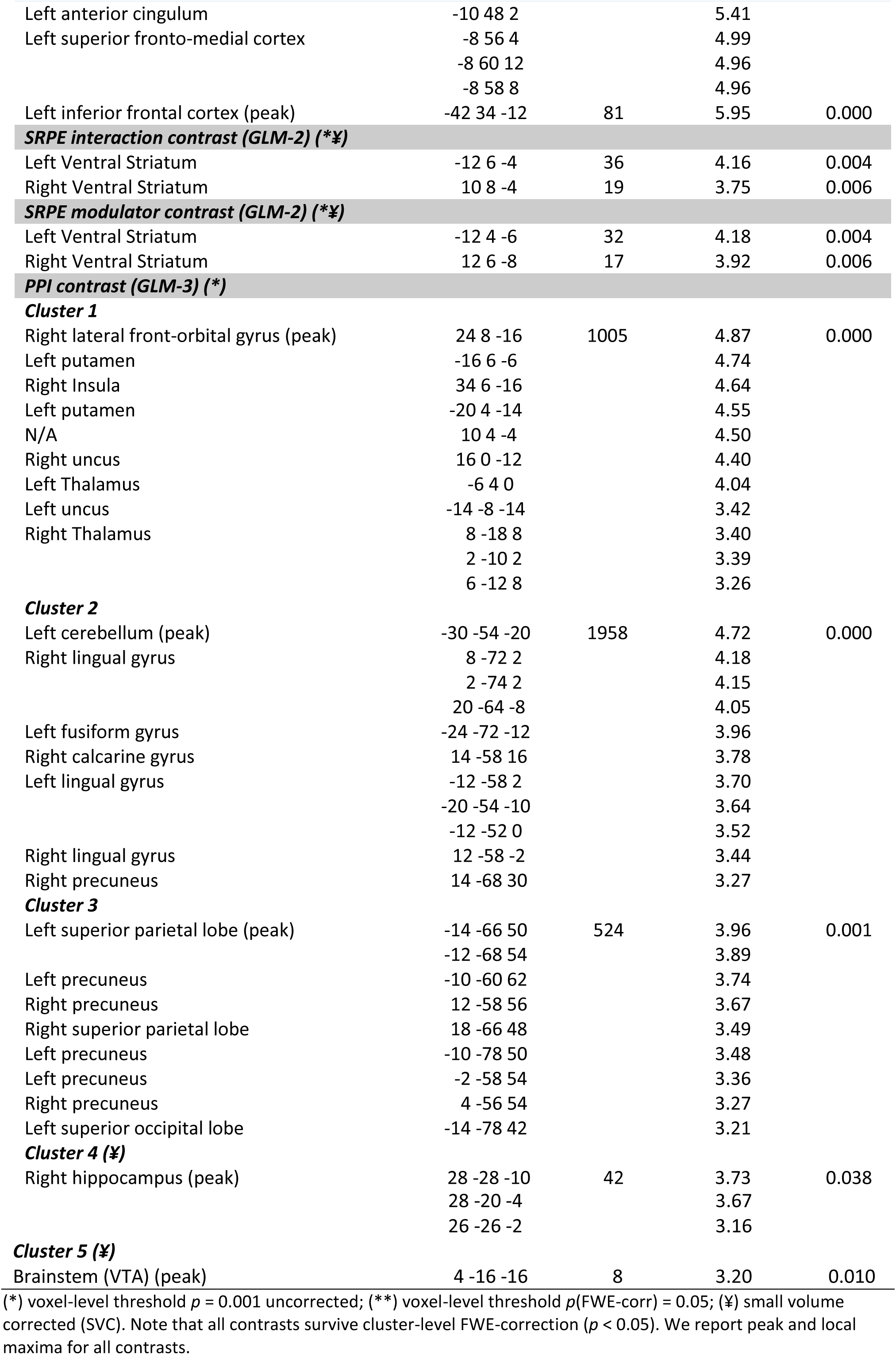
Summary of the activation clusters.

**Table 2.** MNI coordinates of subject-specific face contrast activation maps peak value.

| Subject # | MNI coordinates |  |
| --- | --- | --- |
|  | Left temporal lobe | Right temporal lobe |
| 1 | -46 -54 -16 | 44 -50 -20 |
| 2 | -38 -70 -16 | 46 -48 -22 |
| 3 | -36 -54 -20 | 48 -48 -22 |
| 4 | -48 -54 -18 | 38 -56 -8 |
| 5 | -40 -78 8 | 44 -46 0 |
| 6 | N/A | 52 -76 -4 |
| 7 | -38 -78 4 | 50 -44 4 |
| 8 | -40 -68 -16 | 46 -74 -2 |
| 9 | -44 -46 -14 | 44 -44 -16 |
| 10 | -44 -60 -16 | 34 -56 -12 |
| 11 | -38 -52 -16 | 42 -58 -12 |
| 12 | -46 -50 -22 | 52 -60 -18 |
| 13 | N/A | 40 -56 -16 |
| 14 | -38 -52 -12 | 42 -46 -12 |
| 15 | -46 -60 -14 | 40 -56 -18 |
| 16 | -44 -72 -14 | 36 -54 -16 |
| 17 | -64 -40 -8 | N/A |
| 18 | -38 -44 -26 | 42 -44 -16 |
| 19 | -44 -46 -20 | 48 -48 -24 |
| 20 | -38 -48 -18 | 40 -46 -18 |
| 21 | -42 -64 -14 | 42 -60 -12 |
| 22 | -46 -68 -18 | 42 -54 -18 |
| 23 | -44 -48 -18 | 44 -54 -16 |
| 24 | -36 -48 -16 | 40 -52 -22 |
| 25 | -44 -44 -26 | 42 -68 -16 |
| 26 | -38 -62 -18 | 42 -54 -20 |
| 27 | -46 -70 8 | 46 -52 -16 |
| 28 | N/A | 42 -64 -12 |
| 29 | -38 -80 -12 | 48 -74 -8 |
| 30 | -4 -48 -16 | 40 -42 -18 |

As a first check of our experimental manipulation, we modeled five regressors representing choice feedback for each SRPE value (GLM-1), and contrasted all five regressors (with a contrast vector [1 1 1 1 1]) against baseline (Fig. 2A). If participants were in fact encoding the associations between faces and village names, we expect this contrast to reveal strong hippocampal activation. The choice-feedback contrast indeed revealed robust activation in bilateral hippocampus (left: FWE-*p* < 0.0001, right: FWE-*p* < 0.0001; also violet ROI in Fig. 3B), ventro-medial prefrontal cortex (vmPFC: FWE-*p* < 0.0001), and left inferior frontal cortex (IFC: FWE-*p* < 0.0001).

**Figure 2.**
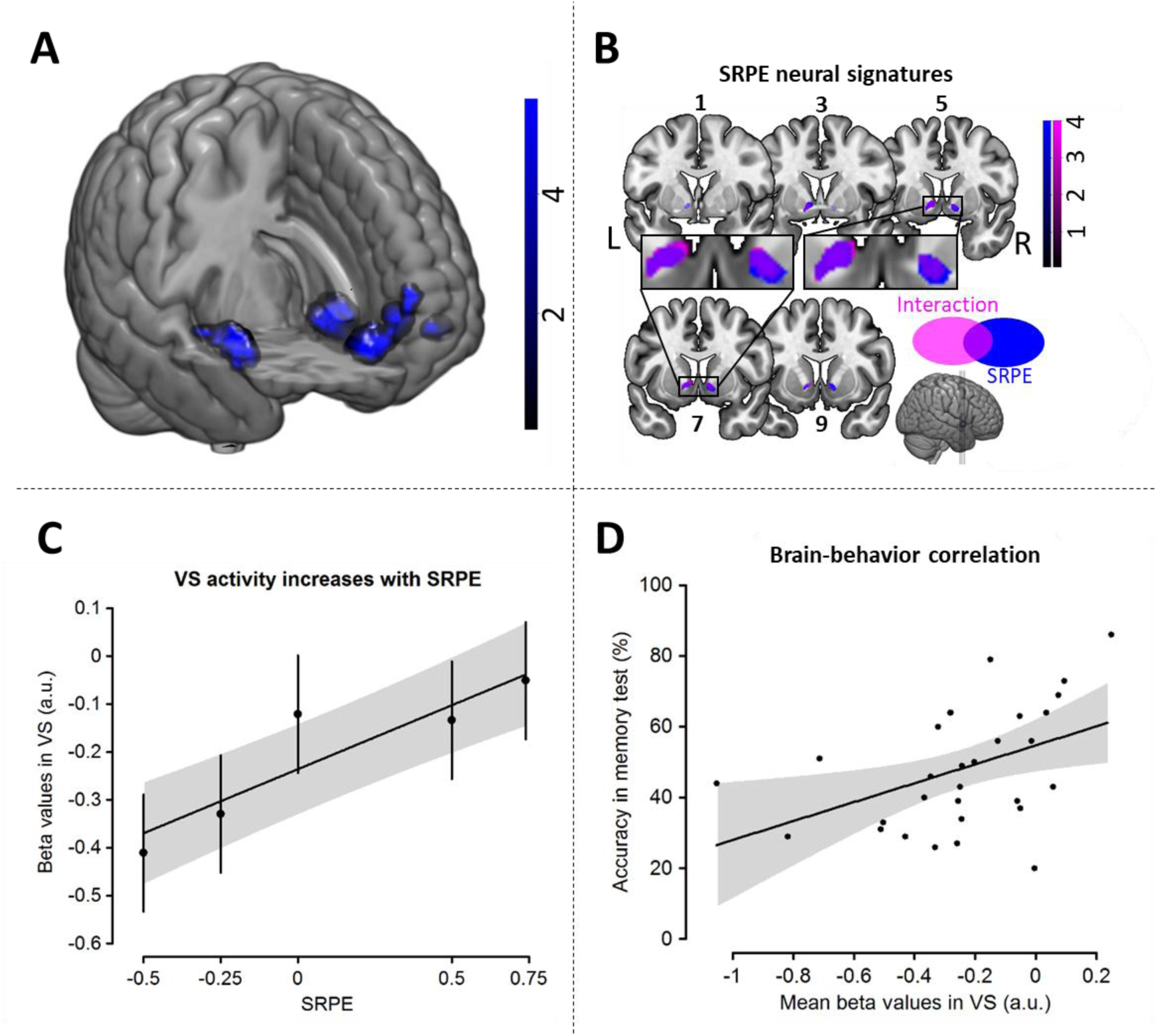
Neural results (see Table 1 for map activation details). **A**.Choice-feedback contrast. The contrast reveals robust bilateral hippocampal (left: z = 5.81, peak [-24, −12, −16]; right: z = 6.19, peak [30, −16, −14]), vmPFC (z = 5.41, peak [-6, 38, −10]), and left IFC (z = 5.95, peak [-42, 34, −12]) activations. **B**. SRPE and SRPE*accuracy interaction contrasts. Bilateral VS cluster (FWE-corrected) maps for SRPE contrast (GLM-1 (blue); left: z = 5.41, peak [-10, 4, −4]; right: z = 5.04, peak [12, 6, −6]), SRPE*subsequent memory accuracy interaction (GLM-2 (violet); left: z = 4.16, peak [-12, 6, −4]; right: z = 3.75, peak [10, 8, −4]). **C**. Exploratory analysis: Positive linear relation between VS activation and SRPE. Mean beta weights extracted from VS (SRPE contrast) for each SRPE value, showing a positive linear relation between SRPE value and VS activation. **D**. Brain-behavior correlation. Positive correlation between mean beta weights extracted from VS and accuracy on the memory test (each dot represents a participant). Numbers above/below brain slices represent y-dimension MNI coordinates. Color bars indicate z-scores. Error bars indicate 95% confidence interval.

**Figure 3.**
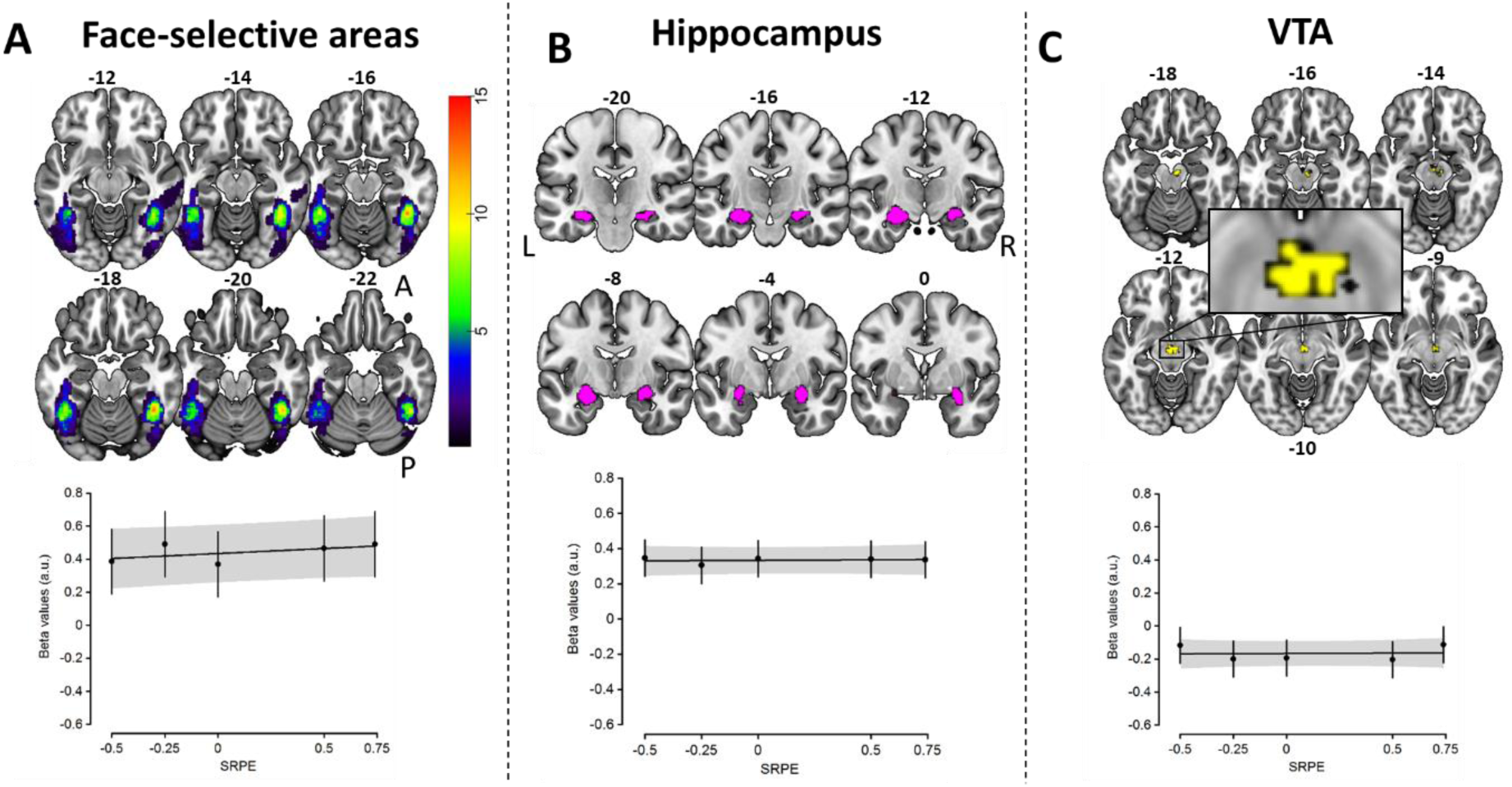
SRPE activation in ROIs. **A**. Face-selective areas. Top: Overlap of subject-specific face contrast activation maps in the inferior temporal lobes. Color bar indicates the number of subjects displaying activation in each voxel; numbers on top of slices indicate z-dimension MNI coordinates. Bottom: No linear relationship was observed between SRPE values and mean activation extracted from FSAs. **B**. Hippocampus. Top: Bilateral hippocampus ROI (violet) extracted from the choice-feedback contrast (Fig. 2A). Numbers on top of slices indicate y-dimension MNI coordinates. Bottom: No linear relationship was observed between SRPE values and mean activation extracted from the hippocampus. **C**. VTA. Top: VTA ROI (yellow) extracted from Neurosynth (see Materials and Methods). Numbers above/under slices indicate z-dimension MNI coordinates. Bottom: No linear relationship was observed between SRPE values and mean activation extracted from VTA.

We then investigated which brain areas encode SRPEs. Based on GLM-1 we tested, at the individual subject level, for a mean-centered SRPE contrast across the five regressors of interest (i.e., the choice-feedback events associated with our five SRPE values). This contrast would hence identify SRPE-sensitive brain areas with increased activity as SRPE value increases. The SRPE contrast revealed robust bilateral VS activations (blue activation in Fig. 2B and Table 1; left VS: FWE-*p* = 0.001; right VS: FWE-*p* = 0.002), suggesting a crucial role of the VS in computing SRPE, in accordance with earlier work (Hyman, Malenka, & Nestler, 2006; Lisman, Grace, & Duzel, 2011; Pine et al., 2018; Scimeca et al., 2016). Notably, no other brain area was identified with this contrast.

To further explore this pattern, we extracted the beta weights from VS for each SRPE level and subject. They displayed a positive linear relationship with SRPE values, *χ*^*2*^(1, *N* = 30) = 40.02, *p* = 2.5e-10 (Fig. 2C). We observed a main effect of reward (*χ*^*2*^(1, *N* = 30) = 51.4, *p* = 1.2e-12; i.e., rewarded trials beta weights have higher values compared with unrewarded trials beta weights). We also observed a main effect of number of options (*t*(29) = 7.05, *p* = 9.279e-08); 4-options trials beta weights have higher values compared with 2-options trials. To further certify that the number of options contributed to the SRPE effect, we performed an exploratory analysis. For each participant we averaged the slope for the negative RPEs (i.e., −0.5 to −0.25) with the slope for the positive RPEs (i.e., 0.5 to 0.75), resulting in the average VS activation increase as the number of options is increased from 2 to 4 options (hence controlling for the unbalanced design, see Fig. 7). We compared the slope of this activation increase against 0 (one-sample t-test, two-sided). Results show a significant increase (*t*(29) = 2.51, *p* = 0.018), suggesting an effect of SRPE which is also instigated by the number of options and not reducible to a pure reward effect. A more rigorous test that the SRPE effect is not reducible to a mere reward effect, is given in the next result section.

**Figure 4A.**
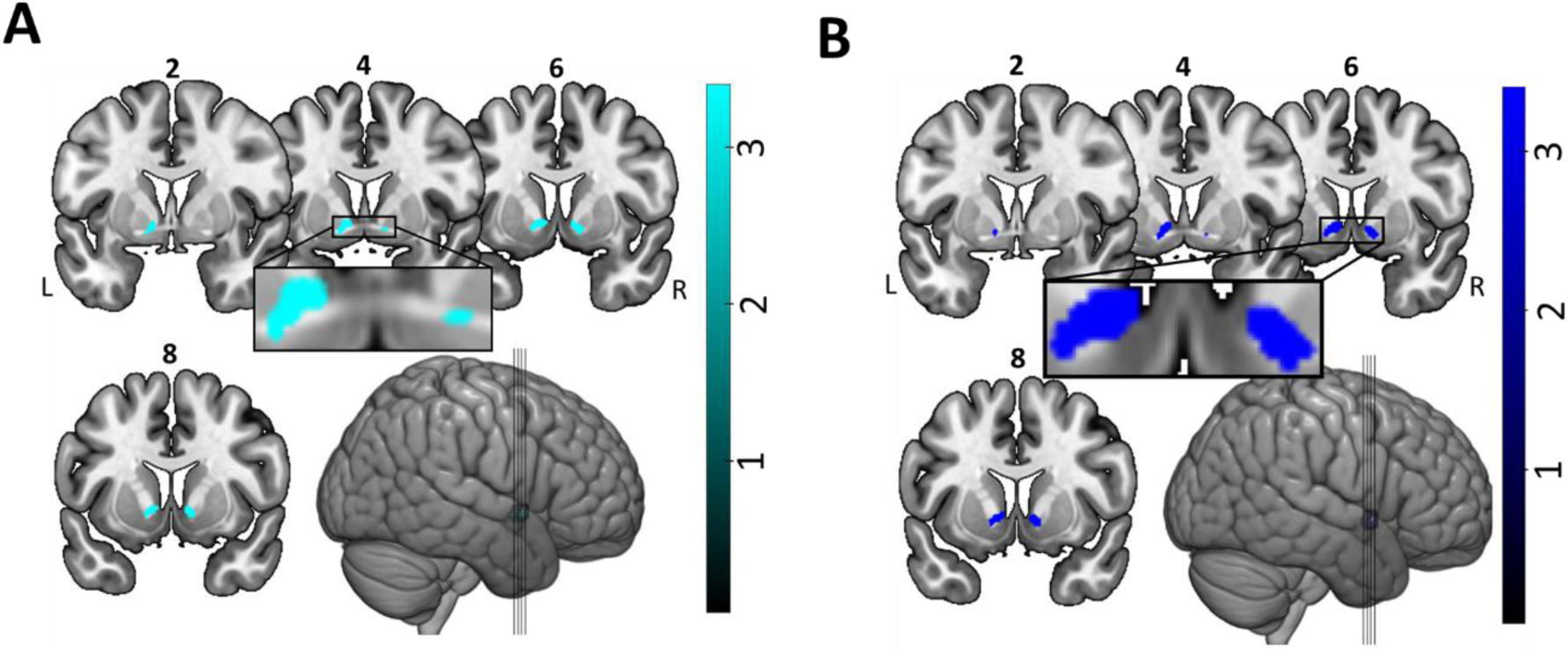
SRPE modulator contrast results. Activation map shows bilateral VS activation (z = 4.18, peak = [-12, 4, −6]; z = 3.92, peak = [12, 6, −8], cluster-level correction). Color bar indicates z-scores; numbers on top of slices indicate y-dimension MNI coordinates. **4B**. Additional analysis on rewarded trials. Activation map shows bilateral VS activation (left: z = 4.51, peak = [-12, 6, −4]; right: z = 4.43, peak = [12, 8, −4], cluster-level correction). Color bar indicates z-scores; numbers on top of slices indicate y-dimension MNI coordinates.

**Figure 5.**
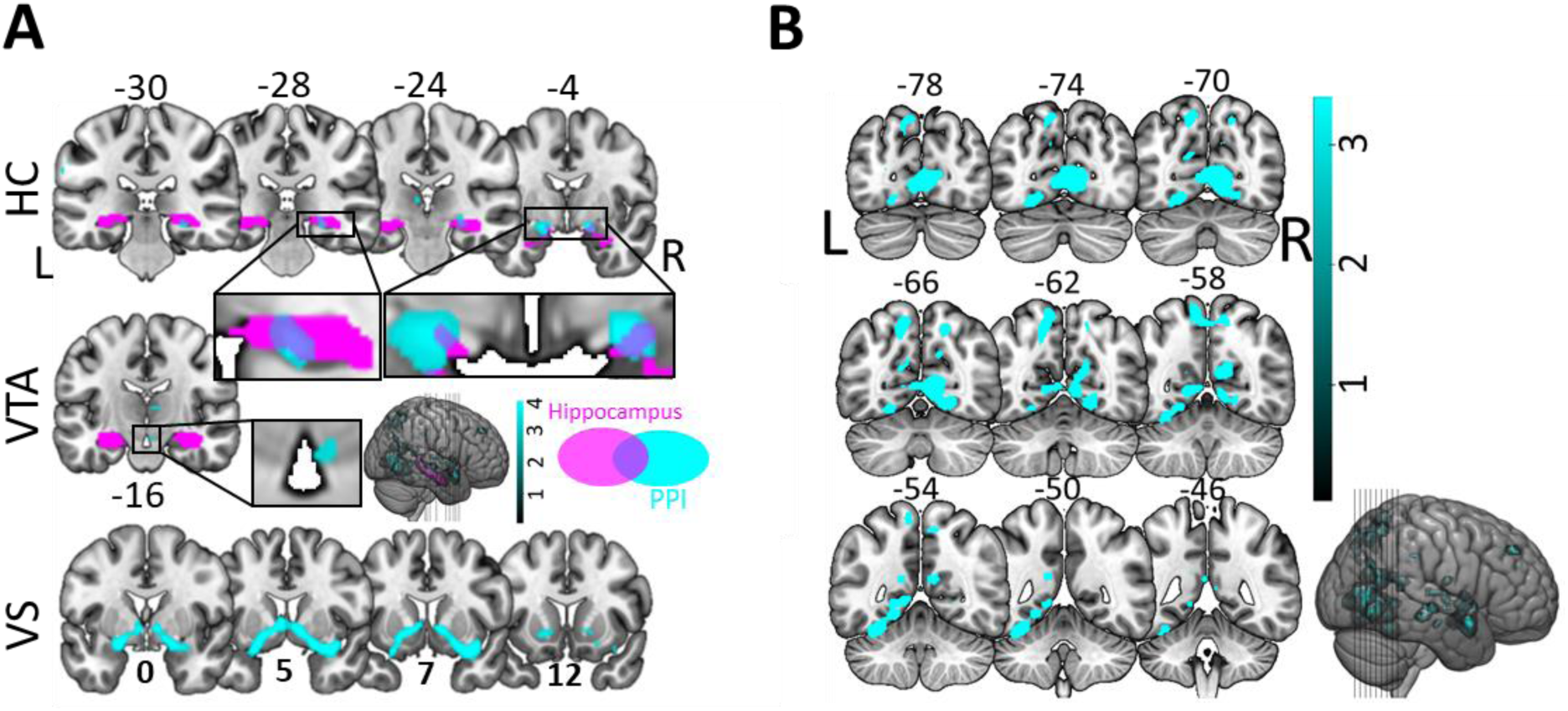
Connectivity results. **A**. Functional connectivity with FSA as a function of SRPE value reveals clusters encompassing hippocampus, VTA and bilateral VS (GLM-3 (cyan); z = 4.87, peak [24, 8, −16]). Purple regions represent bilateral hippocampus from WFU_pickatlas. **B**. Additional connectivity clusters. Activation map shows activation in inferior and medial temporal lobes, as well as superior parietal lobe. Color bar indicates z-scores; numbers on top of slices indicate y-dimension MNI coordinates.

**Figure 6.**
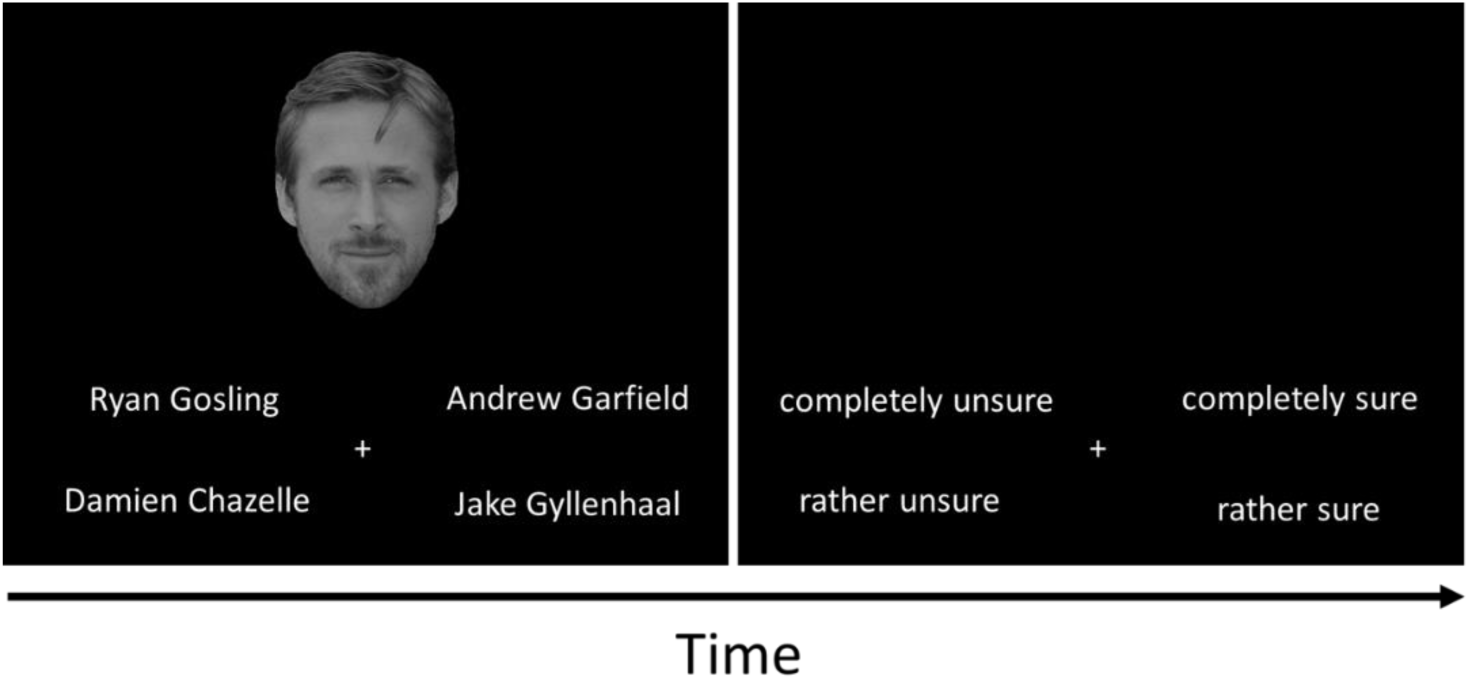
Celebrity knowledge task. Participants were first shown a celebrity face alongside four competing celebrity names. No time limit was imposed to respond with the keyboard (keys “d”, “c”, “n” and “j”, respectively for top left, bottom left, bottom right and top right names). Participants then (using the same keys) indicated their choice certainty.

**Figure 7.**
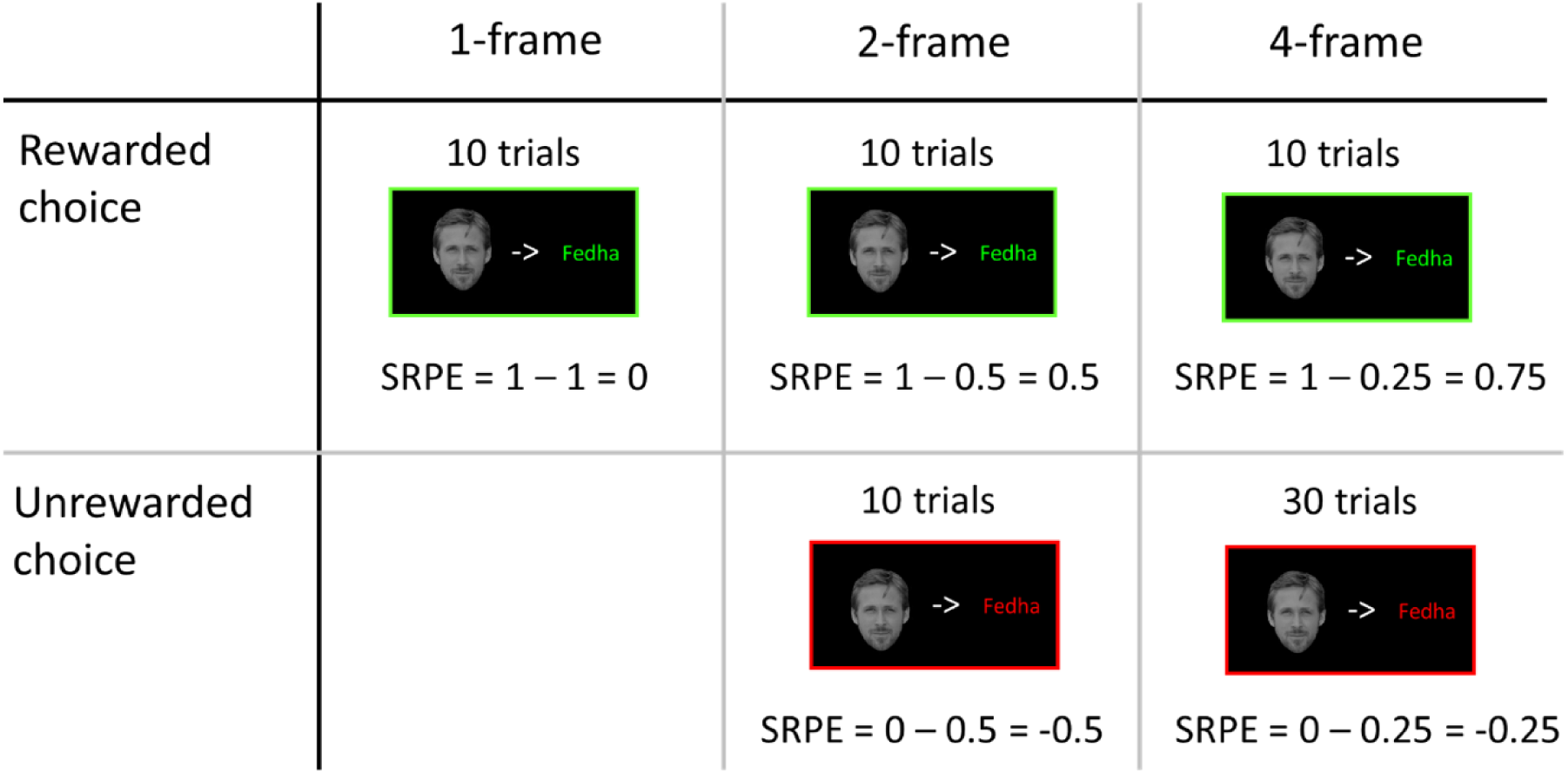
Trial distribution: Number of trials and associated SRPEs.

We further explored whether the linear effect of SRPE on activation was present in our other ROIs. We did not observe such an effect, neither in face-selective areas (FSA, see below), hippocampus, nor VTA (all *p* > 0.16; Fig. 3). Note that we further tested whether specific brain areas would encode for an unsigned RPE (URPE) effect (see Materials and Methods). No brain area was active based on the URPE contrast.

### Striatal SRPEs predict memory accuracy on a trial-by-trial basis

RL theory predicts that steeper SRPE slopes on a trial-to-trial basis would improve subsequent memory. To test this, in a new GLM (GLM-2) we added 4 parametric modulators to a choice feedback regressor (see Materials and Methods). The first modulator indicated whether the encoding phase of that trial led to a correct (1) or wrong (0) recognition at the subsequent memory test (termed recognition accuracy modulator). The second modulator consisted of the SRPE value for the trial at hand; its value could be - 0.5, −0.25, 0, 0.5, or 0.75 (termed SRPE modulator). The third modulator was the SRPE*subsequent memory accuracy interaction (termed the SRPE interaction modulator). Testing for this third modulator identifies brain areas with a stronger SRPE-dependent linear increase driving episodic memory when recognition is correct (relative to wrong) at the subsequent memory test. We added a fourth modulator composed of the interaction between reward (i.e., rewarded (1) or unrewarded (0) choice) and subsequent memory accuracy; allowing us to control for a pure reward*subsequent memory accuracy interaction effect.

The SRPE interaction modulator revealed bilateral activation of the VS (Fig. 2A, violet activation map; left: FWE-*p* = 0.004; right: FWE-*p* = 0.006). This result suggests that VS activation is crucial for successful recognition. Furthermore, the SRPE modulator revealed bilateral VS activation (Fig. 4A, cyan activation map; left: FWE-*p* = 0.004, right: FWE-*p* = 0.006), thereby confirming results from GLM-1. Testing for other modulator contrasts revealed no activation surviving the statistical threshold (whole-brain or with small volume correction (SVC)).

As an additional demonstration that the effect of striatal SRPE value on subsequent memory accuracy is not a pure reward effect, we repeated the GLM-2 analysis only for rewarded trials (i.e., with 3 reward levels 0, .5, and .75). This additional analysis again confirmed our second hypothesis; we observed bilateral activation of the VS (Fig. 4B, blue activation map; left: FWE-*p* = 0.004; right: FWE-*p* = 0.006).

### Functional connectivity between stimulus-processing areas and hippocampus, VTA, and VS depends on SRPE value

Next, we tested if encoding of episodic memory is reflected by increased functional connectivity between stimulus-processing areas, i.e., subject-specific FSAs (Fig. 3A), and hippocampus, VTA and VS, as a function of SRPE value. We thus carried out a psychophysiological interaction (PPI) analysis (Gitelman, Penny, Ashburner, & Friston, 2003) (GLM-3). PPI allows to reveal which areas show an increase in functional connectivity with the seed (i.e., FSAs) as a function of SRPE values. Subject-specific FSAs were revealed using a functional localizer task (see Materials and Methods). Fig. 3A shows the across-subject overlap image statistic displaying where and to what extent FSAs overlap with one another, revealing bilateral activation maps in the inferior temporal lobes. Table 2 reports peak activation of face contrast activation maps for each participant.

The PPI analysis revealed three clusters surviving whole-brain analysis (Fig. 5; FWE-corrected *p* < 0.05, cluster-level). Cluster 1 (Fig. 5A) encompasses activation in bilateral VS, extending to bilateral (anterior) hippocampus, and right amygdala. Cluster 2 (Fig. 5B) displays activation in inferior and medial temporal lobes. Cluster 3 (Fig. 5B) shows activation in superior parietal lobes. Two additional clusters survived theoretically-motivated SVC. Hippocampus (cluster 4) has been involved in previous episodic memory connectivity results and was therefore a theoretically predicted area (Davidow et al., 2016). Cluster 4 displays activation along right hippocampus (Fig. 5A), and was small volume corrected using bilateral WFU pickatlas hippocampus ROIs. Further, VTA (cluster 5) is a small size nucleus generally involved in learning (Bunzeck & Düzel, 2006; Worsley et al., 1996). Cluster 5 displays activation in ventral tegmental area (Fig. 5A), and was small volume corrected using a 6 mm sphere around its peak value. Overall, our PPI results demonstrate that functional connectivity between a stimulus-processing area (FSA) and a network that presumably supports RL of episodic memory (VS, hippocampus, and VTA; (Lisman et al., 2011; Watabe-Uchida et al., 2017)), increases as a function of SRPE. Finally, when taking our other three ROIs as seeds, no significant results were observed.

### Ventral striatum activation predicts overall memory performance

Finally, we tested whether VS activation would be predictive of overall memory performance across subjects. We therefore extracted mean beta weights from the VS (defined from the results of the abovementioned SRPE contrast) to correlate them with performance on the memory test. Our results showed a positive correlation between VS mean beta weights and the percentage of correct recognitions in the memory test (*r* = 0.44; *p* = 0.014; Fig. 2D).

## Discussion

By means of our novel variable-choice task (De Loof et al., 2018; Ergo et al., 2019), we experimentally manipulated SRPE. Using fMRI, we confirmed key RL theory predictions regarding episodic-memory formation. First, we replicated the behavioral effect of signed reward prediction errors (SRPEs) on memory: Stimuli associated to higher SRPE values induce better recognition test performance. Additionally, we confirmed that SRPEs are encoded in the ventral striatum (VS). Second, we revealed a trial-by-trial correlation between striatal SRPE value and subsequent episodic memory. Third, by using localizable task-relevant stimuli (i.e. face-selective area (FSA)), we further showed that connectivity between task-relevant areas and VS, VTA, and hippocampus depends on SRPE. Fourth, we showed that activity in VS correlates (across participants) with overall memory accuracy.

### Signed reward prediction errors influence encoding of episodic information

Our behavioral results replicate earlier work showing that SRPEs drive episodic learning. A stronger better-than-expected reward signal, is associated with better performance at the recognition test (De Loof et al., 2018; Jang et al., 2019). We must consider whether our experimental manipulation confounds the SRPE effect with time-on-task. Importantly, in the 1-frame condition, participants have more time to encode the stimulus associations, given that they already know the correct association at variable-choice onset (i.e., they have 4 more seconds to encode the association compared with the other two conditions). Thus, under a time on task hypothesis, participants would have better memory in the 1-frame condition. Yet, our behavioral data (as earlier ones, see De Loof et al., 2018) show the opposite pattern, ruling out a time on task account.

In our experimental design, the RPE and the to-be-learned face-name association temporally overlapped. Ergo et al. (2019) observed the same behavioral pattern when the feedback was presented prior to the to-be-learned association, demonstrating that a strict temporal overlap is not crucial. It is likely, however, that there is a temporal gradient such that an RPE too far removed from the memorandum leads to diminishing effects. In particular, (Braun et al., 2018) manipulated memorandum – reward temporal (and spatial) distance, and demonstrated such a (temporal and/or spatial) gradient. We predict a similar gradient for RPE effects, although, as the Ergo et al. (2019) data suggest, such a gradient may not be very steep.

In most studies, just a single reward (or RPE) event occurs on each trial. Instead, a recent study manipulated several types of RPE in each trial (Jang et al., 2019). In this study, participants observed the value associated with a specific upcoming gamble (leading to a value-based RPE). Next, they were shown an object (belonging to one of two categories) associated with a specific reward probability (leading to a probability-based RPE). Participants chose to gamble the previously cued value or not (by either selecting or passing on the object), with the instruction that a failed gamble would lead to a negative earning. After their choice, they received reward feedback (leading to a feedback-based RPE). They demonstrated that only the RPE arriving at the same time as the memorandum, yielded an effect on subsequent memory. This finding suggests that if one RPE occurs simultaneously with the memorandum, there is little room for other RPEs to still exert such an effect.

In addition to the time scale of the RPE-memorandum difference, there is the time scale at which participants are tested. Several studies have shown the temporal robustness of the SRPE effect on episodic memory formation, lasting up to 24h (Davidow et al., 2016; De Loof et al., 2018; Jang et al., 2019) or even a week (Pine et al., 2018). This demonstrates that the RPE effect operates at time scales that can support learning and memory in the real world.

### Striatal SRPEs predict subsequent memory

Rewarded versus neutral to-be-memorized items have been found to activate (dopaminergic) midbrain, hippocampus, and striatal activation (Wittmann et al., 2005). Moreover, rewarded items elicit stronger activation in the ventral striatum (as well as in dopaminergic midbrain and hippocampus), when subsequently remembered versus forgotten (Adcock et al., 2006). These findings support a role of reward processing in episodic learning. The effect of reward on memory is thought to be due to dopaminergic signaling in the striatum, presumably leading to enhanced long-term potentiation (LTP) of the to-be-remembered stimuli (Lisman et al., 2011).

Striatal activation associated to episodic learning does not necessarily need to be linked to explicit reward. For instance, successfully learning the meaning of new words caused striatal activation, similar to that of a reward gambling task (Ripollés et al., 2014). In addition, activity in bilateral ventral striatum increased when a previously learned word was recognized as being correctly used (i.e., made semantic sense) in a subsequent sentence, suggesting that self-monitoring of successful novel word learning may be associated with potential reward-related signals (Ripollés et al., 2016). Interestingly, novel word learning performance is enhanced when participants are given levodopa (a dopaminergic precursor) compared with risperidone (a dopamine antagonist) (Ripollés et al., 2018).

In addition, SRPE has been related to subsequent memory. In a recent study, participants learnt episodic information and were tested on their recognition in two subsequent tests, respectively 2 days (Test 1) and 7 days (Test 2) after learning. VS activity (measured in Test 1) was negatively correlated with choice-confidence ratings in Test 2, specifically when participants transitioned from an incorrect (Test 1) to a correct response (Test 2) (Pine et al., 2018). Another study reported a negative relationship between striatal RPEs and subsequent memory performance (Wimmer, Braun, Daw, & Shohamy, 2014), seemingly in contradiction with our current data and with RL theory. The discrepancy between this and the current study may relate to how RPEs are calculated in each. Specifically, in one of the studies (Wimmer et al., 2014), subjects must track reward probability, thus imposing a challenging dual task on top of the memory task. In such case, trials with a strong RPE signal may signify that subjects were more attentive to the reward tracking task, at a cost for the memory task. Instead, our novel variable-choice paradigm imposes no dual-task requirements: The reward probability was explicitly and clearly presented on each single trial. In such case, we indeed observed a positive relationship between ventral striatal RPEs and subsequent memory encoding.

In addition to the effect of SRPE during learning, as our data show, striatal RPEs may also play an important role during retrieval. For instance, during an old-new item decision task, biased positive feedback (i.e., overall more positive feedback regardless of decision accuracy) induces a shift in the decision criterion (towards a more lenient criterion) (Scimeca et al., 2016). Thus, striatal RPEs play a role in the strategies used to make memory decisions. Together with ours, these results suggest that RL may support episodic memory both during encoding and retrieval.

### SRPE value modulates connectivity strength between stimulus-relevant areas and the episodic memory network

Several studies have demonstrated changes in connectivity profiles within (parts of) the episodic learning network prior to, during, and after encoding the to-be-remembered stimuli.

For instance, when participants are cued that an upcoming to-be-remembered item is associated with a high reward, connectivity between VTA and hippocampus is increased during the cue interval (i.e. prior encoding) for subsequently remembered versus forgotten items (Adcock et al., 2006).

Other studies showed that connectivity strength between hippocampus and striatum (putamen) is larger for rewarded (correct) than for unrewarded (incorrect) items. This result was observed at the moment of incidental item encoding, and only present in adolescents but not in adults, suggesting a learning advantage to increased reward sensitivity in adolescents.

An important aspect of our experimental design relies on the type of stimuli used. Specifically, participants had to remember face-word associations. Therefore, by using an independent face-selective area localizer, we were able to test whether changes in connectivity profiles could take place between stimulus-processing areas and a previously reported RPE-based learning network consisting of VTA, VS and hippocampus. We could test this at the moment of stimulus association encoding. The results confirmed our prediction: Connectivity between face-selective areas (our stimulus-processing area) and the RPE-based episodic learning network increases with SRPE values, i.e. the stronger the experienced SRPE, the stronger the coupling with the aforementioned areas. These data are in line with the neoHebbian framework (Lisman et al., 2011), suggesting that synaptic learning between two neurons is a (multiplicative) function of pre- and postsynaptic activity, modulated by a dopaminergic reward (here, SRPE) signal.

Changes in connectivity profiles have already been observed post-stimulus encoding. Pre-versus post-learning changes in the VTA-hippocampus resting-state functional connectivity (RSFC) predicted reward-related memory recognition advantages (Gruber, Ritchey, Wang, Doss, & Ranganath, 2016). In the same vein, differences in connectivity strength prior to and post-stimulus encoding have been shown to be modulated by reward context. In particular, it has been shown that the connectivity between category-selective areas (i.e. FFA and parahippocampal place area (PPA)) and hippocampus is enhanced when reward is high (Murty, Tompary, Adcock, & Davachi, 2017).

Although RPE-dependent changes in connectivity profiles have been demonstrated, the question of how RPEs increase connectivity and thereby improve memory remains. One possibility may be that RPEs enhance theta phase synchronization. In line with such a view, experimentally induced theta synchronization between visual and auditory modalities improved multimodal stimulus memories (Clouter, Shapiro, & Hanslmayr, 2017), and theta phase synchronization in such a paradigm (as measured via EEG) predicts memory performance on a trial-by-trial basis (Wang, Clouter, Chen, Shapiro, & Hanslmayr, 2018). Therefore, theta synchronization between relevant brain areas may play the role of efficiently cementing memory information.

### Conclusions

We manipulated SRPE in an episodic associative learning task. We replicated previous work showing the behavioral effect of SRPE on subsequent memory, i.e. high SRPE values lead to enhanced subsequent memory. Furthermore, we observed that SRPEs, encoded in VS, predict both, across subjects and on trial-to-trial basis, subsequent memory accuracy. Finally, we demonstrated that connectivity strength between stimulus-processing areas and VS, VTA, and hippocampus, is modulated by SRPE values. Therefore, we suggest that episodic memory encoding is guided by an RPE-based neural (RL) mechanism, as is also the case in procedural learning (Schultz, Dayan, & Montague, 1997). In episodic memory, bilateral VS relays VTA/SN-computed RPEs towards stimulus-processing areas and hippocampus, thereby increasing functional connectivity between hippocampus and cortical areas.

## Materials and Methods

In this section, we provide all technical, methodological, and analytical details. We start by describing participant demographics, followed by a full description of the experimental design. We then successively explain behavioral and fMRI data acquisition and analyses. All tasks and analysis codes, as well as face, word, and house stimuli used in our experiment are available on OSF (https://osf.io/6vkwm/?view_only=8b1364c4cb6b41d4b0e96ad615827d33).

### Participants

Thirty right-handed participants (26 females, mean age = 22, s.d. = 6.62; 4 males, mean age = 30, s.d. = 11.15) with normal vision participated in this study approved by the local ethics committee (Ghent University-Hospital (UZ Gent), Ghent, Belgium). Participants received monetary compensation (30 euro) and provided written informed consent prior to the experiment. Our sample size was motivated by previous studies in our lab, robustly showing the sought for behavioral effect, using the same number of participants (De Loof et al., 2018; Ergo et al., 2019).

### Experimental design

Participants underwent four tasks (total duration ∼80 min) in the following order: celebrity knowledge task (outside the scanner, ∼15 min), variable-choice and functional localizer tasks (inside the scanner, ∼50 min), and memory test (outside the scanner, ∼15 min). All tasks were programmed with PsychoPy2.

#### Celebrity knowledge task

In the celebrity knowledge task, participants were shown a celebrity face alongside four potential celebrity names. Participants had to select the correct celebrity name by pressing the keys “d”, “c”, “n” or “j” to respectively select the name in the top left, bottom left, bottom right, or top right corner of the display (Fig. 6). Following their choice, participants gave a certainty rating on a 4-point scale ranging from “completely unsure”, “rather unsure”, “rather sure” to “completely sure”. They gave their answer using the same keys as above. Participants had no time restriction on these choices. The celebrity knowledge task was performed in one block of 140 trials. Once completed, we selected a subset of 70 accurately recognized faces, which had been rated as “completely sure”. When participants did not reach that level of accuracy, a random draw of faces was selected to complete the subset of 70 trials. This subset of 70 faces composed the stimulus set subsequently used in the variable-choice task for that subject (see below). We chose known celebrities because these stimuli can be verbalized and, hence, increase the possibility of learning the association between faces and village names (see below). On top of this, we randomly selected six of the remaining faces for the variable-choice task training, and a set of 60 of the remaining faces (not used in the variable-choice task) for the functional localizer task. This task was run on a Dell Latitude E5550 laptop.

#### Variable-choice task

Participants underwent the variable-choice task in the scanner (Fig. 1A). They first observed a fixation cross (0.5 sec), followed by the presentation of a celebrity face on top of the screen together with four village pseudo-names. After exploring the display for four seconds, either 1, 2 or 4 names were framed. The participant’s task was to guess which village name was associated with the celebrity face. Participants were constrained to choose between the framed names. By varying the number of framed village names, we were able to manipulate the signed reward prediction error (SRPE) on each trial. We can compute the SRPE as *r – p*, where *r* is the observed reward (1 and 0 for correct and incorrect guesses, respectively), and *p* is the probability of making a correct guess. This probability is 1, 0.5 or 0.25, respectively for the 1-, 2- or 4-frame conditions. Hence, SRPE could take on the values −0.5, −0.25, 0, 0.5, 0.75. Participants made their choice (no time restrictions) using Cedrus Lumina LS-Pair MRI compatible ergonomic response pads. To select the upper left, bottom left, bottom right or top right option, participants respectively pressed with their left middle finger, left index, right index or right middle finger. Following their choice, a fixation cross was shown for a jittered amount of time, drawn from a Poisson distribution (λ = 4) truncated between 1 and 7 seconds. Subsequently, during the choice-feedback (5 sec), participants were shown the celebrity face associated to the correct village name, either framed in green (if they guessed correctly) or in red (otherwise). Choice-feedback was followed by another jittered fixation cross, drawn from a Poisson distribution (λ = 4) truncated between 1 and 4 seconds. The trial ended by a monetary update (2 sec) indicating the money earned for that trial (0.70 euro for correct guesses and 0 euro otherwise), as well as a total tally. Both fixation cross jitters were optimized to maximize design efficiency using the DesignDiagnostics toolbox (https://montilab.psych.ucla.edu/fmri-wiki/).

The variable-choice task was performed in one block of 70 trials. Figure 7 displays the distribution of trials in each condition. The 1-frame, 2-frame, and 4-frame conditions were respectively composed of 10, 20, and 40 trials of which 10 trials (in each case) led to a correct guess (i.e., rewarded choice).

Note that prior to the variable-choice task, participants underwent six training trials outside of the scanner; two of each condition (i.e., 1-, 2-, and 4-frame) leading to correct and incorrect guesses (except for the 1-frame condition always leading to correct guesses). Therefore, participants had experienced all the SRPEs prior to performing the task inside the scanner. The training phase was run on a Dell Latitude E5550 laptop, and participants gave their response with the “d”, “c”, “n” or “j” key (as in the celebrity knowledge task).

#### Functional localizer task

Immediately following the variable-choice task, participants underwent the functional localizer task. In this task, participants alternatingly observed a centrally presented celebrity face or a house (1.5 sec) and a fixation cross (0.5 sec). The task of the participants was to respond with the right index finger, whenever the presentation of a face/house would repeat (i.e., 1-back task; Fig. 8). Note that contrasting blocks of a 1-back task on face versus house stimuli has been proven efficient to functionally reveal fusiform face area (FFA) activation (Berman et al., 2010) (see below). House pictures were taken from earlier work (Schiffer, Muller, Yeung, & Waszak, 2014) and faces were randomly selected from the subset of 60 faces that was not presented during the variable-choice task. Participants performed 16 blocks (8 blocks with faces and 8 with houses, in random order) of 18 trials. Each stimulus had a 0.2 probability of repeating itself (but could not repeat itself twice).

**Figure 8.**
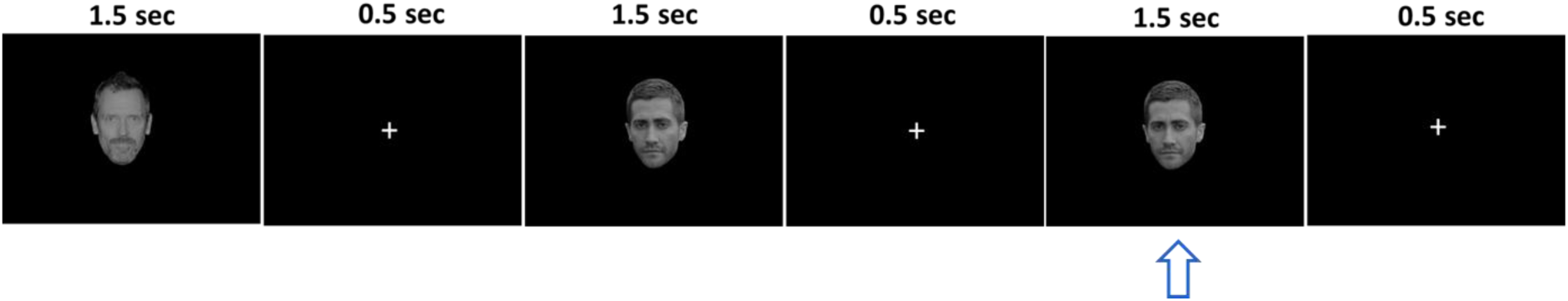
Functional localizer task. Participants observed a series of celebrity faces (or houses) and had to press with their right index finger whenever a face (or house) was repeated (blue arrow).

#### Memory test

Upon completion of the functional localizer task, participants performed the memory test (Fig. 9). During this test, participants again observed the 70 faces alongside the same four competing village names (shuffled relative to their previous positions on the display during the variable-choice task). In order to minimize primacy or recency effects, we chunked the 70 trials presented during the variable-choice task in chunks of 10 trials; the first 10 trials forming chunk 1, the following 10 trials forming chunk 2, and so on. The trials of each chunk were randomly shuffled, and the chunks were represented in sequential order (from chunk 1 to 7). As in the celebrity knowledge task, participants pressed keys “d”, “c”, “n”, or “j” to respectively select the village names at the top left, bottom left, bottom right or top right; and were instructed to provide a certainty rating on their choice after each trial. We imposed no time restrictions on either task.

**Figure 9.**
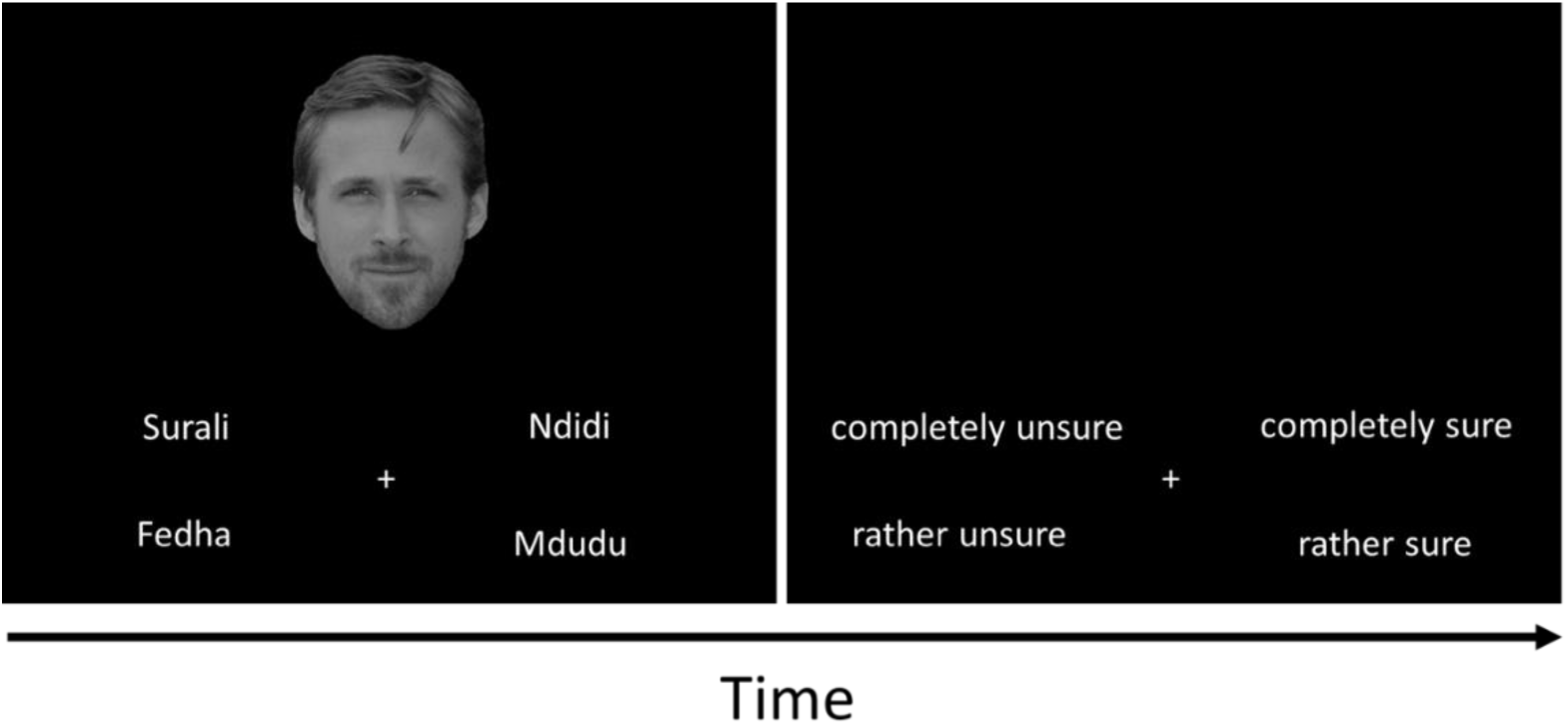
Trial structure of the memory test. Participants observed a celebrity face alongside four village names. Using the same keys as in the celebrity knowledge task, participants recalled the association previously learned during the variable-choice task. Subsequent to their choice, participants rated their choice certainty (using the same keys).

### Behavioral data analysis

To test the behavioral effect of SRPE on recognition performance, we applied a (generalized) linear mixed effects model with a random intercept per subject and centered predictors. To combine the behavioral with the fMRI results, trial characteristics (e.g., accuracy and neural activation) were averaged per SRPE level for each participant. We report the *χ*^*2*^ statistics from the ANOVA Type III tests. All analyses were performed in R.

### fMRI data acquisition and analysis

#### fMRI data acquisition

fMRI data were collected with a 3T Magnetom Trio MRI scanner system (Siemens Medical Systems, Erlangen, Germany), with a thirty-two-channel radio-frequency head coil. A 3D high-resolution anatomical image was obtained using a T1-weighted MPRAGE sequence (TR = 2200 ms, TE = 2.51 ms, TI = 900 ms, FA = 8°, FOV: 250 × 250 mm, matrix size: 176 × 256 × 256, interpolated voxel size: 0.9 × 0.9 × 0.9 mm). In addition, we acquired a field map per participant, to correct for magnetic field inhomogeneities (TR = 520 ms, TE1 = 4.92 ms, TE2 = 7.38 ms, image matrix = 80 × 80 x 50, FOV = 192 mm, FA = 60°, slice thickness = 2.5 mm, voxel size = 2.5 × 2.5 × 2.5 mm, distance factor = 20%). Whole brain functional images were acquired using a T2*-weighted EPI sequence (TR = 1730 ms, TE = 30 ms, FA = 66°, multiband acceleration factor = 2, matrix size: 50 × 84 × 84; voxel size: 2.5 × 2.5 × 2.5 mm).

#### fMRI data analysis

SPM 12 (https://www.fil.ion.ucl.ac.uk/spm/software/spm12/; Matlab R2016b) was used for preprocessing and fMRI data analysis. For each participant, the first five volumes were discarded to avoid transient spin saturations effects, and BIDS (https://bids.neuroimaging.io/) formatted raw data was defaced, realigned and unwarped, slice time corrected, normalized, and smoothed using a Gaussian kernel of 8 mm full width at half maximum (FWHM). We performed co-registration of the structural image with the mean realigned functional image. Additionally, time series were inspected for excessive movement using the Artifact Detection Toolbox (ART; https://www.nitrc.org/projects/artifact_detect), allowing us to regress out functional volumes displaying excessive motion/spikes (see below).

Each participant’s data was modeled with a general linear model (GLM) using an event-related design. To test our different empirical predictions, we performed three distinct GLMs. Common to all GLMs, regressors were convolved with the canonical HRF, and the cut-off period for high-pass filtering was 128 seconds. Six movement parameters, derived from spatial realignment, as well as spikes detected using ART were included as covariates of non-interest. First-level summary statistic images were entered in a second-level analysis in which subjects were treated as random effects (RFX). Our neural predictions are based on choice-feedback related activity (Fig. 1A). We created three general linear models (GLM) to test them. GLM-1 tested for the SRPE effect, expected to be encoded in VS (Pine et al., 2018), and whether VS activity correlates (across subjects) with subsequent episodic memory. GLM-2, tested for the interaction between striatal SRPE value and subsequent episodic memory accuracy. GLM-3 tested for the PPI between SRPE (psychological factor) and subject-specific face-selective areas (FSA) time series (physiological factor). Below we describe in detail our three GLMs.

First, as a check of our experimental manipulation and to reveal the area encoding SRPE, we constructed GLM-1. In GLM-1, we modeled the reading phase (i.e., the joint onset of faces and names) as a boxcar function (4 sec), and the choice (i.e., response press) and the monetary update as delta functions. Of main interest, we added five regressors of choice-feedback (boxcar, 5 sec) respectively for each SRPE value (i.e., −0.5, −0.25, 0, 0.5, 0.75). To check our experimental manipulation, we tested for an overall effect of choice-feedback, termed choice-feedback contrast, by contrasting all five SRPE regressors of interest (see above; with a contrast vector [1 1 1 1 1]) to baseline. Given that participants were asked to memorize the association between faces and words, we expected this contrast to reveal hippocampal activation. To reveal the area encoding SRPE, we then tested, at the individual level, for a mean-centered SRPE contrast [-0.6 −0.35 −0.1 0.40 0.65] across the five regressors of interest (i.e., the choice-feedback events associated with our five SRPE values). Any area sensitive to this contrast would display increased activity as SRPE value increases, and would therefore encode for SRPE. Given previous results (Davidow et al., 2016; Pine et al., 2018), we predicted that ventral striatum (VS) would encode for SRPE (although statistical correction was at whole-brain level). Furthermore, to test whether VS activation predicts overall subsequent memory performance, we extracted choice-feedback locked mean beta weights from the VS ROI, again defined using the group-level SRPE contrast (FWE-corrected, *p* < 0.05; blue activation map, Fig. 2B). We then correlated mean beta weights and subsequent memory accuracy (in %) across subjects. We expected a positive linear relationship between activity in VS and memory accuracy. Furthermore, we carried out two exploratory analyses based on GLM-1. In a first exploratory analysis, we assessed the SRPE contrast by extracting mean beta weights for each of the five SRPE regressors in four distinct empirically derived ROIs (Pine et al., 2018); all four ROIs were active in earlier episodic memory studies (Davidow et al., 2016; Ripollés et al., 2016). First, we focused on the VS, which was defined using the group-level SRPE contrast (FWE-corrected, *p* < 0.05; blue activation map, Fig. 2B). Second, we focused on the VTA (Cohen, Haesler, Vong, Lowell, & Uchida, 2012). Given that no contrast revealed a clear VTA activation, we used Neurosynth (http://neurosynth.org/) to define our ROI. We searched for the term “Ventral Tegmental” and applied a threshold of 18 on z-scores of the activation map, yielding our VTA ROI. Third, hippocampus was defined using the group-level results of the choice-feedback contrast (see above). Fourth, we considered the face-selective areas (FSA) with the face contrast (see GLM-4 below). In a second exploratory analysis, we tested for an unsigned reward prediction error (URPE) over the five RPE levels in GLM-1. The URPE values are calculated by taking the absolute value of the SRPE values followed by centering, to form the URPE contrast [0.1 −0.15 −0.4 0.1 0.35]. Such contrast would reveal any potential areas encoding for surprise, irrespective of whether surprise has a positive or negative valence.

Second, to test whether SRPE value in VS correlates with memory on a trial-by-trial basis, we constructed GLM-2. GLM-2 was similar to GLM-1 except as follows. The choice feedback was modeled as one regressor of interest (instead of five); irrespective of SRPE value. We further added four parametric modulators and turned off orthogonalization. The first modulator indicated whether the encoding phase of that trial led to a correct (1) or wrong (0) recognition at the subsequent memory test (termed subsequent memory contrast). The second modulator consisted of the SRPE value for the specific trial at hand; its value could be −0.5, −0.25, 0, 0.5, or 0.75 (termed SRPE modulator contrast). The third modulator was the SRPE*subsequent memory interaction. Testing for this third modulator identifies brain areas with a stronger SRPE-dependent linear increase driving episodic memory when recognition is correct (relative to wrong) at the memory test (termed the SRPE interaction contrast). We added a fourth modulator composed of the interaction between reward (i.e., rewarded (1) or unrewarded (0) choice) and subsequent memory accuracy; allowing us to control for a reward*subsequent memory accuracy interaction effect. To exclude any interpretation of the SRPE interaction contrast in terms of a pure reward effect, we repeated the previous analysis including only rewarded trials.

Third, we carried out a psychophysiological interaction (PPI) analysis (Gitelman et al., 2003) (GLM-3). PPI allows to reveal which other areas show an increase in functional connectivity with the seed stimulus-relevant area, as a function of SRPE values. In GLM-3, three regressors were added (on top of the nuisance regressors described in GLM-1). The first regressor consisted of the BOLD signal extracted from a 3 mm sphere (Davidow et al., 2016) around the peak value from subject-specific functionally localized face-selective area (FSA seed, see below; see Table 2 for MNI coordinates of subject-specific peak values). The second regressor is the psychological vector consisting of the SRPE contrast as described in GLM-1. The third regressor is the seed-by-condition interaction. Areas that show a linear increase in coupling with the seed as a function of SRPE value will be identified by a significant seed-by-condition interaction regressor. To gain a deeper understanding of the functional connectivity between our ROIs, we also performed the PPI analysis using our hippocampus, VTA and VS ROIs (defined above) as seeds of interest.

Our functional localizer task was modeled using a block design (GLM-4). Each block of the 1-back task was modeled using a boxcar function over the entire block duration. Nuisance regressors were modeled as in GLM-1. To reveal subject-dependent functional FSA, we contrasted face vs. house blocks (*p* < 0.0001, uncorrected). Using the resulting statistical maps and MRIcron, we created an overlap image statistic displaying where and to what extent FSAs overlap with one another.

## Acknowledgements

The current work was supported by grant G0F3818N (EOS grant FWO / FNRS). C.B.C is supported by FWO grant #12O7719N. K.E. is supported by FWO grant #1153420N. We thank Carlos González-García and Clay Holroyd for helpful comments.

## References

Adcock, R. A., Thangavel, A., Whitfield-Gabrieli, S., Knutson, B., & Gabrieli, J. D. E. (2006). Reward-Motivated Learning: Mesolimbic Activation Precedes Memory Formation. Neuron, 50(3), 507–517. https://doi.org/10.1016/j.neuron.2006.03.036

Berman, M. G., Park, J., Gonzalez, R., Polk, T. A., Gehrke, A., Knaffla, S., & Jonides, J. (2010). Evaluating functional localizers: The case of the FFA. NeuroImage, 50(1), 56–71. https://doi.org/10.1016/j.neuroimage.2009.12.024

Braun, E. K., Wimmer, G. E., & Shohamy, D. (2018). Retroactive and graded prioritization of memory by reward. Nature Communications, 9(1), 4886. https://doi.org/10.1038/s41467-018-07280-0

Bunzeck, N., & Düzel, E. (2006). Absolute Coding of Stimulus Novelty in the Human Substantia Nigra/VTA. Neuron, 51(3), 369–379. https://doi.org/10.1016/j.neuron.2006.06.021

Clouter, A., Shapiro, K. L., & Hanslmayr, S. (2017). Theta Phase Synchronization Is the Glue that Binds Human Associative Memory. Current Biology, 27(20), 3143-3148.e6. https://doi.org/10.1016/j.cub.2017.09.001

Cohen, J. Y., Haesler, S., Vong, L., Lowell, B. B., & Uchida, N. (2012). Neuron-type-specific signals for reward and punishment in the ventral tegmental area. Nature, 482(7383), 85–88. https://doi.org/10.1038/nature10754

Davidow, J. Y., Foerde, K., Galván, A., & Shohamy, D. (2016). An Upside to Reward Sensitivity: The Hippocampus Supports Enhanced Reinforcement Learning in Adolescence. Neuron, 92(1), 93–99. https://doi.org/10.1016/j.neuron.2016.08.031

De Loof, E., Ergo, K., Naert, L., Janssens, C., Talsma, D., Van Opstal, F., & Verguts, T. (2018). Signed reward prediction errors drive declarative learning. PLOS ONE, 13(1), e0189212. https://doi.org/10.1371/journal.pone.0189212

Ergo, K., De Loof, E., Janssens, C., & Verguts, T. (2019). Oscillatory signatures of reward prediction errors in declarative learning. NeuroImage, 186(June 2018), 137–145. https://doi.org/10.1016/j.neuroimage.2018.10.083

Gitelman, D. R., Penny, W. D., Ashburner, J., & Friston, K. J. (2003). Modeling regional and psychophysiologic interactions in fMRI: The importance of hemodynamic deconvolution. NeuroImage, 19, 200–207. https://doi.org/10.1016/S1053-8119(03)00058-2

Gruber, M. J., Ritchey, M., Wang, S. F., Doss, M. K., & Ranganath, C. (2016). Post-learning Hippocampal Dynamics Promote Preferential Retention of Rewarding Events. Neuron, 89(5), 1110–1120. https://doi.org/10.1016/j.neuron.2016.01.017

Hyman, S. E., Malenka, R. C., & Nestler, E. J. (2006). NEURAL MECHANISMS OF ADDICTION: The Role of Reward-Related Learning and Memory. Annual Review of Neuroscience, 29(1), 565–598. https://doi.org/10.1146/annurev.neuro.29.051605.113009

Jang, A. I., Nassar, M. R., Dillon, D. G., & Frank, M. J. (2019). Positive reward prediction errors during decision-making strengthen memory encoding. Nature Human Behaviour, 327445. https://doi.org/10.1038/s41562-019-0597-3

Lisman, J., Grace, A. A., & Duzel, E. (2011). A neoHebbian framework for episodic memory; role of dopamine-dependent late LTP. Trends in Neurosciences, 34(10), 536–547. https://doi.org/10.1016/j.tins.2011.07.006

Miendlarzewska, E. A., Bavelier, D., & Schwartz, S. (2016). Influence of reward motivation on human declarative memory. Neuroscience and Biobehavioral Reviews, 61, 156–176. https://doi.org/10.1016/j.neubiorev.2015.11.015

Murty, V. P., Tompary, A., Adcock, R. A., & Davachi, L. (2017). Selectivity in Postencoding Connectivity with High-Level Visual Cortex Is Associated with Reward-Motivated Memory. The Journal of Neuroscience, 37(3), 537–545. https://doi.org/10.1523/JNEUROSCI.4032-15.2016

Pine, A., Sadeh, N., Ben-Yakov, A., Dudai, Y., & Mendelsohn, A. (2018). Knowledge acquisition is governed by striatal prediction errors. Nature Communications, 9(1), 1–14. https://doi.org/10.1038/s41467-018-03992-5

Ripollés, P., Ferreri, L., Mas-Herrero, E., Alicart, H., Gómez-Andrés, A., Marco-Pallares, J., … Rodriguez-Fornells, A. (2018). Intrinsically regulated learning is modulated by synaptic dopamine signaling. ELife, 7, 1–23. https://doi.org/10.7554/eLife.38113

Ripollés, P., Marco-Pallarés, J., Alicart, H., Tempelmann, C., Rodríguez-Fornells, A., & Noesselt, T. (2016). Intrinsic monitoring of learning success facilitates memory encoding via the activation of the SN/VTA-Hippocampal loop. ELife, 5, 1–35. https://doi.org/10.7554/eLife.17441

Ripollés, P., Marco-Pallarés, J., Hielscher, U., Mestres-Missé, A., Tempelmann, C., Heinze, H.-J., … Noesselt, T. (2014). The Role of Reward in Word Learning and Its Implications for Language Acquisition. Current Biology, 24(21), 2606–2611. https://doi.org/10.1016/j.cub.2014.09.044

Schiffer, A.-M., Muller, T., Yeung, N., & Waszak, F. (2014). Reward Activates Stimulus-Specific and Task-Dependent Representations in Visual Association Cortices. Journal of Neuroscience, 34(47), 15610–15620. https://doi.org/10.1523/JNEUROSCI.1640-14.2014

Schultz, W., Dayan, P., & Montague, P. R. (1997). A neural substrate of prediction and reward. Science, 275(5306), 1593–1599. https://doi.org/10.1126/science.275.5306.1593

Scimeca, J. M., Katzman, P. L., & Badre, D. (2016). Striatal prediction errors support dynamic control of declarative memory decisions. Nature Communications, 7. https://doi.org/10.1038/ncomms13061

Shneyer, A., & Mendelsohn, A. (2018). Previously rewarding environments enhance incidental memory formation. Learning and Memory, 25(11), 569–573. https://doi.org/10.1101/lm.047886.118

Sutton, R., & Barto, A. (2018). Reinforcement Learning: An Introduction.

Tulving, E. (1993). What Is Episodic Memory? Current Directions in Psychological Science, 2(3), 67–70. https://doi.org/10.1111/1467-8721.ep10770899

Wang, D., Clouter, A., Chen, Q., Shapiro, K. L., & Hanslmayr, S. (2018). Single-trial phase entrainment of theta oscillations in sensory regions predicts human associative memory performance. Journal of Neuroscience, 38(28), 6299–6309. https://doi.org/10.1523/JNEUROSCI.0349-18.2018

Watabe-Uchida, M., Eshel, N., & Uchida, N. (2017). Neural Circuitry of Reward Prediction Error. Annual Review of Neuroscience, 40(1), 373–394. https://doi.org/10.1146/annurev-neuro-072116-031109

Wimmer, G. E., Braun, E. K., Daw, N. D., & Shohamy, D. (2014). Episodic Memory Encoding Interferes with Reward Learning and Decreases Striatal Prediction Errors. Journal of Neuroscience, 34(45), 14901–14912. https://doi.org/10.1523/jneurosci.0204-14.2014

Wittmann, B. C., Schott, B. H., Guderian, S., Frey, J. U., Heinze, H. J., & Düzel, E. (2005). Reward-related fMRI activation of dopaminergic midbrain is associated with enhanced hippocampus-dependent long-term memory formation. Neuron, 45(3), 459–467. https://doi.org/10.1016/j.neuron.2005.01.010

Worsley, K. J., Marrett, S., Neelin, P., Vandal, A. C., Friston, K. J., & Evans, A. C. (1996). A unified statistical approach for determining significant signals in images of cerebral activation. Human Brain Mapping, 4(1), 58–73. https://doi.org/10.1002/(SICI)1097-0193(1996)4:1<58::AID-HBM4>3.0.CO;2-O

